# Site-specific DNA demethylation during spermatogenesis precedes nucleosome retention in mouse sperm

**DOI:** 10.1101/2025.01.10.632457

**Authors:** So Maezawa, Masashi Yukawa, Akihiko Sakashita, Yuto Yamada, Tomoko Sagi, Artem Barski, Satoshi H. Namekawa

## Abstract

DNA methylation patterns are inherited from the parental germline to the embryo. In mature sperm, the sites of unmethylated DNA are tightly coupled to sites of histone retention at gene regulatory elements that are implicated in paternal epigenetic inheritance. The timing and mechanism of site-specific DNA demethylation in the male germline remains unknown. Here, we perform genome-wide profiling of DNA methylation during spermatogenesis by capturing methylated DNA through interaction with a methyl-DNA binding protein domain (MBD). Our data demonstrate a site-specific change in DNA methylation during the mitosis-to-meiosis transition. Importantly, demethylation at these genomic sites during this transition precedes nucleosome retention in spermatozoa. These results suggest that site-specific DNA demethylation during the mitosis-to-meiosis transition of spermatogenesis contributes to the establishment of sperm epigenetic states associated with embryonic gene expression. We therefore propose DNA demethylation during spermatogenesis as a novel phase of epigenetic reprogramming that contributes to embryonic gene regulation.

## Introduction

Methylation at the 5-carbon position of cytosine (5-methylcytosine: 5mC) at CpG dinucleotides in DNA is dynamically reprogrammed in the parental germline and contributes to epigenetic inheritance from gametes to embryos (Greenberg & Bourc’his, 2019). In mammalian sperm, the vast majority of histones are replaced by protamines, but sites of unmethylated DNA are tightly coupled to sites of histone retention at gene regulatory elements (Erkek et al., 2013). These gene regulatory elements are associated with the embryonic gene expression program after fertilization (Brykczynska et al., 2010; Carone et al., 2014; Erkek et al., 2013; Hammoud et al., 2009; Jung et al., 2019; Jung et al., 2017; Yamaguchi et al., 2018; Yoshida et al., 2018) and therefore contribute to paternal epigenetic inheritance (Gaspa-Toneu & Peters, 2023; Lismer & Kimmins, 2023).

However, it remains unknown when the hypomethylated regions associated with histone retention in spermatozoa are established in the male germline.

The sites of histone retention exhibit features of bivalent genomic domains. These domains are characterized by concomitant enrichment of repressive Polycomb Repressive Complex 2 (PRC2)-mediated trimethylation of histone H3 at lysine 27 (H3K27me3) and active di/trimethylation of H3 at lysine 4 (H3K4me2/3) (Brykczynska et al., 2010; Erkek et al., 2013; Hammoud et al., 2014; Hammoud et al., 2009; Lesch et al., 2013; Sachs et al., 2013). Recent studies have demonstrated that alteration of the H3K27me3 and H3K4me3 levels in sperm leads to changes in gene expression in the next generation (Lesch et al., 2019; Lismer et al., 2021; Sakashita et al., 2023; Siklenka et al., 2015).

In the male germline, DNA methylation is largely reprogrammed in primordial germ cells. It globally increases in embryonic prospermatogonia and reaches a level similar to that in spermatozoa in postnatal spermatogonia (∼80% of all CpG sites) (Hammoud et al., 2014; Kobayashi et al., 2013; Kubo et al., 2015; Seisenberger et al., 2012). In prospermatogonia, DNA methylation outside of CpG islands, i.e. clusters of CpG dinucleotides that are constitutively hypomethylated, is established through the action of the histone methyltransferase NSD1, the de novo DNA methyltransferase DNMT3A and its catalytically inactive cofactor DNMT3L (Kaneda et al., 2004; Kato et al., 2007; Shirane et al., 2020). Evolutionarily young transposable elements are recognized by the PIWI-interacting RNA (piRNA) pathway and methylated by the germline- specific de novo DNA methyltransferase DNMT3C (Aravin et al., 2007; Barau et al., 2016; Jain et al., 2017; Kuramochi-Miyagawa et al., 2008).

Based on the studies of paternal genomic imprinting loci, it has long been thought that DNA methylation is stably maintained throughout spermatogenesis following its acquisition in prospermatogonia (Abramowitz & Bartolomei, 2012; Sasaki & Matsui, 2008; Schaefer et al., 2007). Accumulating evidence suggests a more dynamic nature of DNA methylation during postnatal spermatogenesis. At the postnatal stage, changes in DNA methylation were initially detected in testicular germ cells (Oakes et al., 2007) and during spermatogonial differentiation (Shirakawa et al., 2013). Later studies clarified that, during spermatogenesis, male germ cells undergo a global transient reduction of DNA methylation in early meiotic prophase I (Gaysinskaya et al., 2018; Huang et al., 2023). This process is presumably passive and caused by a delay in establishing maintenance DNA methylation by DNMT1 and its cofactor UHRF1 during premeiotic DNA replication (Gaysinskaya et al., 2018; Huang et al., 2023). Other genome-wide studies have also detected modest changes in DNA methylation during spermatogenesis (Ben Maamar et al., 2022; Chen et al., 2020; Liu et al., 2019; Siebert-Kuss et al., 2024).

However, the extent of site-specific regulation of DNA methylation during postnatal spermatogenesis beyond the global transient reduction remains unclear.

To detect site-specific changes in DNA methylation during spermatogenesis, we performed MethylCap-seq during representative stages of spermatogenesis. This approach employs capture of methylated DNA via the Methyl-CpG-binding domain (MBD) followed by next-generation sequencing (Brinkman et al., 2010). Our analysis shows that a site-specific loss of DNA methylation during the mitosis-to-meiosis transition precedes nucleosome retention sites in mature sperm. Our findings suggest that meiotic DNA demethylation contributes to the establishment of sperm epigenetic states associated with embryonic gene expression.

## Results

### DNA methylation dynamics during spermatogenesis

During mouse spermatogenesis, maintenance DNA methylation is delayed following premeiotic DNA replication, resulting in transient hemimethylation during premeiotic S phase. DNA methylation is then gradually restored during meiotic prophase I (Gaysinskaya et al., 2018; Huang et al., 2023) (Fig. 1A). To evaluate the overall change before and after this transient reduction of DNA methylation, we first quantified the amount of 5mC during spermatogenesis using an enzyme-linked immunosorbent assay (ELISA). Specifically, we analyzed postnatal day 7 (P7) THY1^+^ undifferentiated spermatogonia and KIT^+^ differentiating spermatogonia, both cellular stages prior to the transient reduction. To cover stages of late spermatogenesis after the transient reduction of DNA methylation, we analyzed meiotic pachytene spermatocytes (PS) and postmeiotic round spermatids (RS) from adult testes. We found the 5mC level increased during the initial spermatogonial differentiation (Fig. 1B). After the transient reduction, 5mC levels in PS did not recover to the level of KIT^+^ differentiating spermatogonia, but after meiosis, they slightly increased in RS (Fig. 1B).

**Figure 1:**
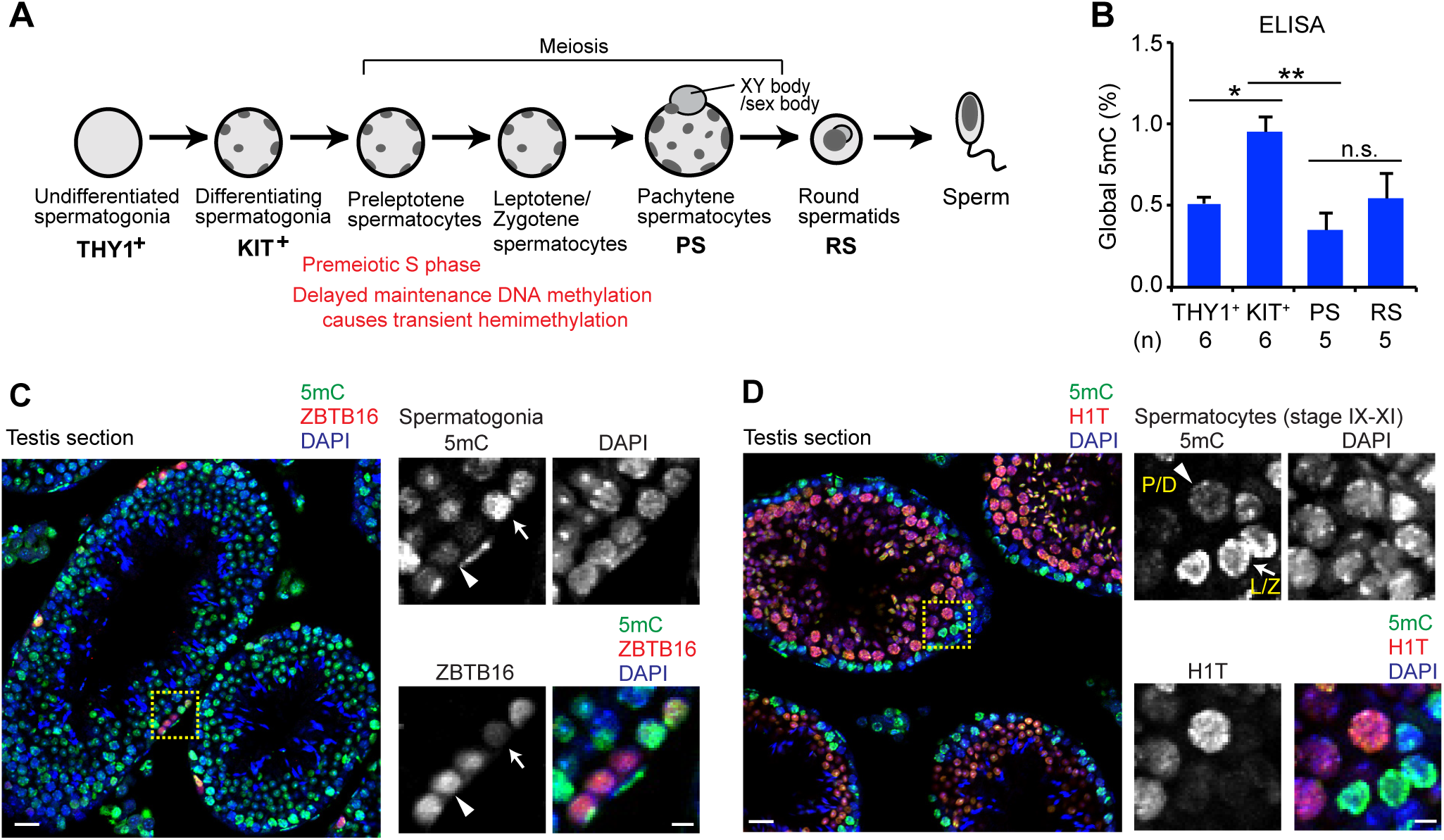
Global characteristics of DNA methylation during spermatogenesis. (A) Schematic of spermatogenesis and the four representative stages analyzed in this study (THY1^+^, KIT^+^, PS, and RS). The previously reported period of delayed maintenance methylation following premeiotic DNA replication, which results in transient hemimethylation, and the subsequent restoration of DNA methylation are indicated. (B) ELISA-based quantification of global 5mC levels during spermatogenesis. The percentage of 5mC in total DNA is shown. Data are presented as mean ± SEM from six biological replicates for THY1-positive and KIT-positive spermatogonia and five biological replicates for PS and RS. Statistical significance was assessed by one-way ANOVA followed by Tukey’s multiple-comparison test. *P < 0.05; **P < 0.01; n.s., not significant. (C, D) Immunostaining of paraffin sections of adult testes with anti-5mC antibody and antibodies against stage-specific markers (ZBTB16 and H1T). Slides were counterstained with DAPI. Images were acquired with a confocal microscope. Regions bordered by dashed yellow squares are magnified in the right panels. Scale bar in the left panel: 20 µm; scale bar in the right panel: 5 µm.

To independently evaluate these observations and further define the timing of DNA methylation dynamics during spermatogenesis, we performed immunohistochemistry on testicular sections using stage-specific germ cell markers. Consistent with the ELISA results, nuclear 5mC signals were higher in KIT-positive differentiating spermatogonia (Fig. S1A) than in ZBTB16 (PLZF)-positive undifferentiated spermatogonia (Fig. 1C). This finding is consistent with the previously reported increase in DNA methylation during spermatogonial differentiation (Shirakawa et al., 2013). Strong 5mC signals were also observed in STRA8^+^ preleptotene spermatocytes and DMC1^+^ leptotene spermatocytes. In contrast, markedly reduced 5mC signals were detected in γH2AX^+^ pachytene spermatocytes and in pachytene/diplotene spermatocytes expressing the testis-specific histone variant H1T, a marker induced after the mid-pachytene stage (Fig. 1D and Fig. S1B). These results demonstrate dynamic changes in overall 5mC levels during spermatogonial differentiation and meiotic progression.

### Detection of site-specific DNA methylation dynamics in spermatogenesis

To investigate locus-specific DNA methylation remodeling underlying the global changes observed in Figure 1, we performed MethylCap-seq (Brinkman et al., 2010), which captures methylated DNA through the methyl-CpG-binding domain. Unlike whole-genome bisulfite sequencing, which does not distinguish between 5mC and 5-hydroxymethylcytosine, MethylCap-seq specifically detects 5mC. While MethylCap-seq does not offer base-pair resolution, it does provide overall profiles of 5mC genome-wide, particularly on CpG-dense regions (Nair et al., 2011). In addition, we reasoned that because MethylCap-seq depends on the recognition of 5mC by MBD, the readout of this method represents the recognition by MBD, which may be relevant to functional aspects of 5mC.

We performed MethylCap-seq in wild-type P7 THY1^+^ and KIT^+^ spermatogonia and adult PS and RS with two biological replicates. We detected approximately 130,000 to 200,000 common peaks genome-wide at each transition (Fig. 2A, Table S1), demonstrating that the overall MethylCap-seq profile largely overlapped between each stage. This observation is in line with the fact that male germ cells acquire global DNA methylation levels similar to spermatozoa before birth. Compared to the number of these common peaks, we detected a relatively small number of unique peaks during spermatogonial differentiation (between THY1^+^ and KIT^+^ spermatogonia) and during the meiosis-to-postmeiosis transition (between PS and RS) (Fig. 2A). In contrast, at the transition from KIT^+^ spermatogonia to PS, more site-specific changes in 5mC were detected compared to the other transitions (15,285 KIT^+^ unique peaks and 20,934 PS unique peaks). The distribution of these unique peaks encompasses various genomic regions, including intergenic regions, introns, exons, promoters, and upstream regions (Fig. 2A, right panel). A representative track view of a ∼ 100-kb genomic region in chromosome 17 shows examples of site-specific 5mC gains and losses at the transition between KIT^+^ spermatogonia and PS, and these patterns are consistent between biological replicates (Fig. 2B). Average tag density analysis revealed that the loss of 5mC at the KIT^+^ to PS transition is pronounced at transcription start sites (TSSs) of late spermatogenesis genes, which are activated at the KIT^+^ to PS transition (Sin et al., 2015) (Fig. 2C). Demethylation at TSSs of late spermatogenesis genes, such as *Spata16* and *Slc22a16*, was confirmed both in the MethylCap-seq data and previously-published WGBS data (Hammoud et al., 2014; Kubo et al., 2015) (Fig. S2A). Consistent with this finding, hypomethylation of spermatogenesis genes was recently reported in a human study (Siebert-Kuss et al., 2024). In addition, we observed extensive demethylation at the protocadherin (*Pcdh*) genes of the *Pcdh*γ locus (Fig. S2B). In contrast, 5mC levels did not change extensively in other classes of genes, such as somatic/progenitor genes that are turned off at the KIT^+^ to PS transition or genes that are constitutively active or inactive during spermatogenesis (Sin et al., 2015) (Fig. 2C). These observations prompted us to examine promoter DNA methylation dynamics in greater detail.

**Figure 2:**
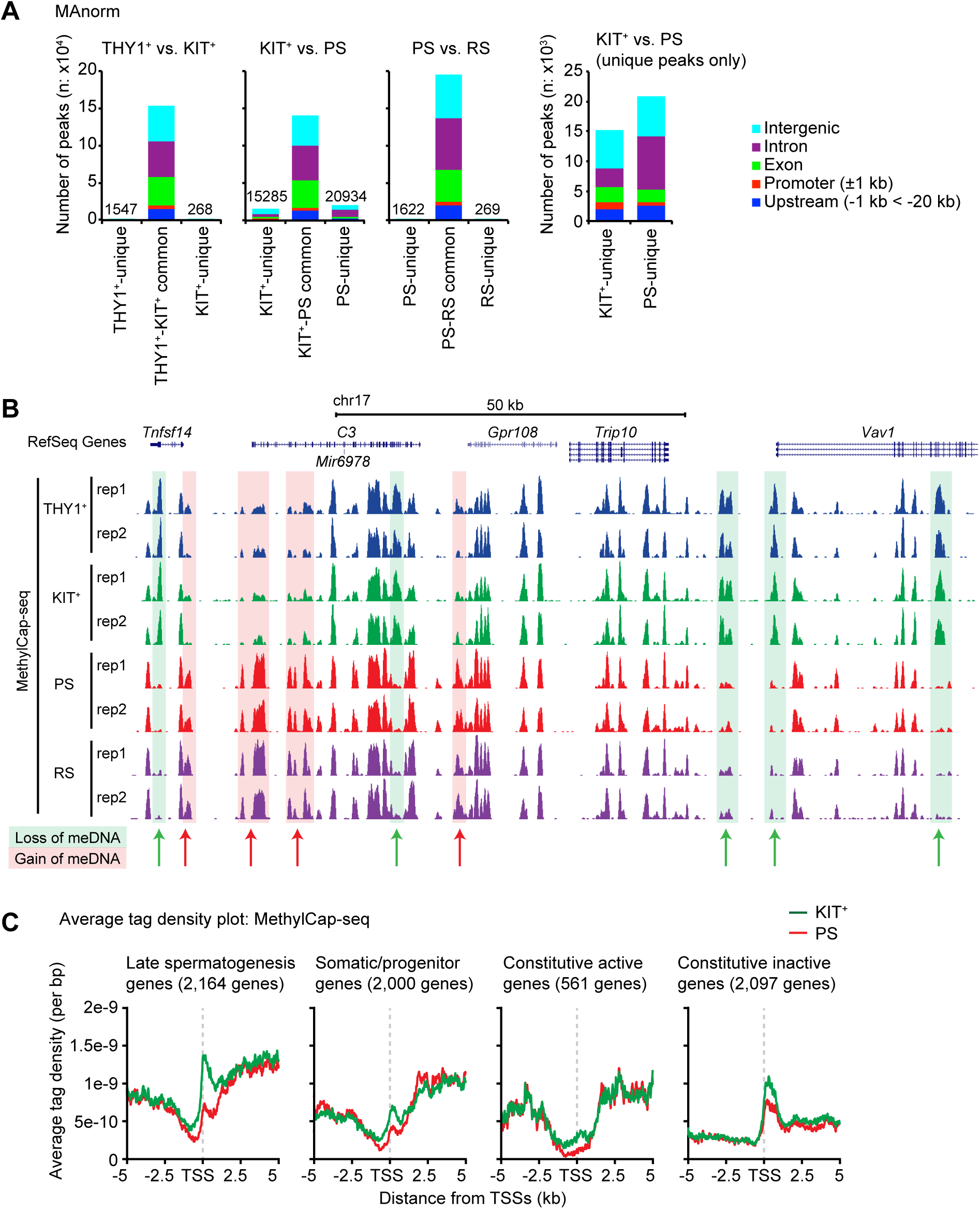
MethylCap-seq analysis during spermatogenesis. (A) Dynamic changes in DNA methylation during spermatogenesis. MAnorm analysis of MethylCap-seq at each transition of spermatogenesis. The genomic distribution of each peak is shown with colored bars. (B) Track view of MethylCap-seq data of a representative genomic region for each stage of spermatogenesis. Two biological replicates are shown. Sites with loss of DNA methylation are underlaid in green and indicated by green arrows; sites with gain of DNA methylation are underlaid in red and indicated by red arrows. (C) Average tag density plots of MethylCap-seq reads in each representative group of genes defined in a previous study (Sin et al., 2015).

### Promoter demethylation accompanies activation of late spermatogenesis genes

To further define the changes in the 5mC level at TSSs during spermatogenesis, we compared 5mC levels around all TSSs between two stages through scatter plots of MethylCap-seq read enrichment. Consistent with the peak analysis, 5mC levels around TSSs between THY1^+^ and KIT^+^ spermatogonia and between PS and RS are highly correlated (Fig. 3A). KIT^+^ and PS showed a decreased correlation compared to the other transitions (Fig. 3A), although it is still high. These data show overall moderate changes in the 5mC levels between the different cellular stages analyzed. Nevertheless, at the KIT^+^ to PS transition, 909 TSSs showed a loss of 5mC levels, while 753 TSSs showed a gain of 5mC levels (Fig. 3B, Table S2).

**Figure 3:**
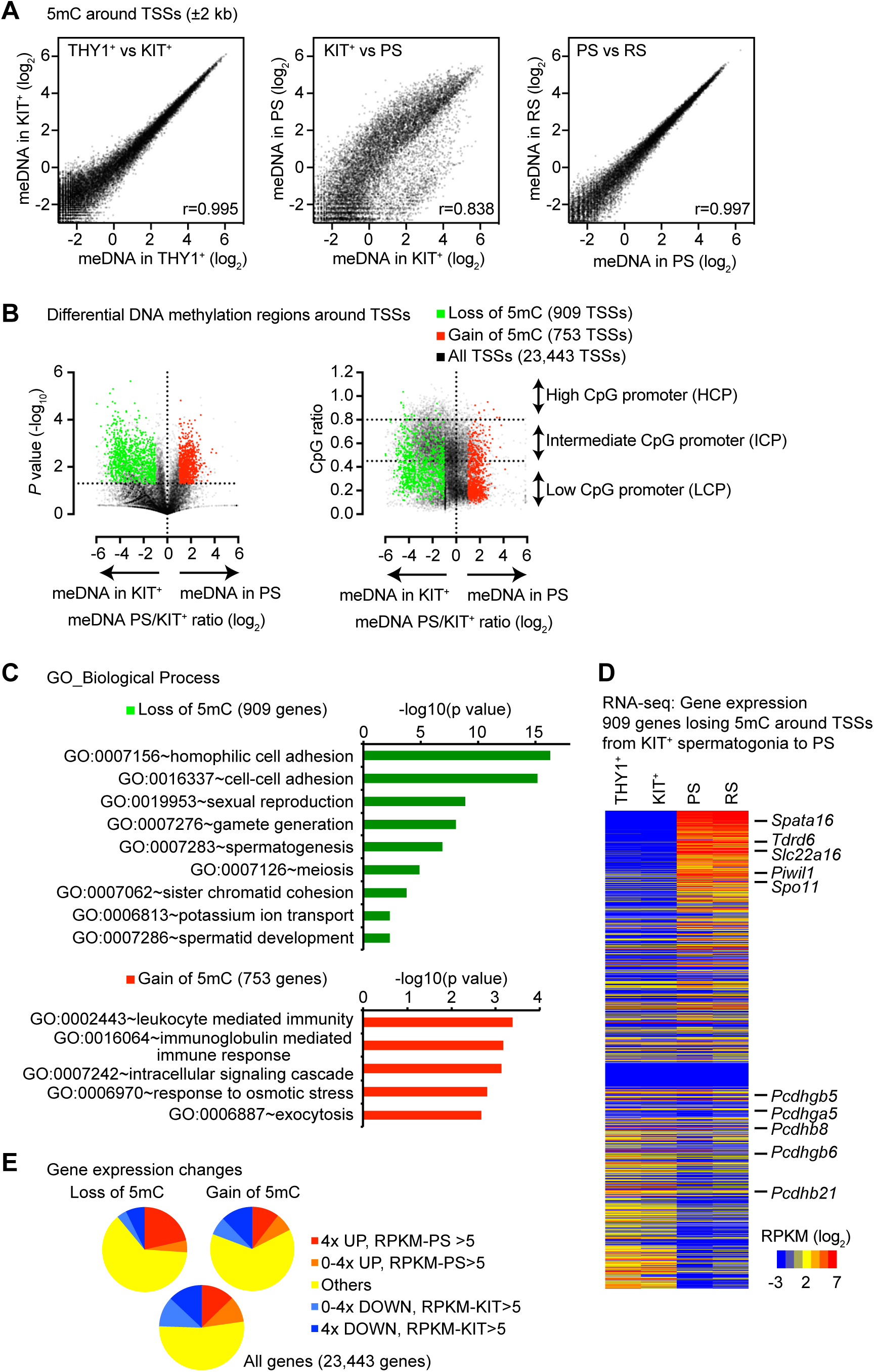
Changes in DNA methylation during spermatogenesis. (A) Enrichment analysis of MethylCap-seq around TSSs (±2 kb) at each transition during spermatogenesis. Pearson correlation values are shown. (B) Volcano plots for the enrichment and P-values of MethylCap-seq enrichment within ±1 kb of TSSs during the transition from KIT^+^ spermatogonia to PS. Peaks were detected by MAnorm. (C) GO enrichment analyses for 909 genes losing 5mC and 753 genes gaining 5mC from KIT^+^ spermatogonia to PS. (D) RNA-seq heatmap for 909 genes losing 5mC from KIT^+^ spermatogonia to PS. (E) Pie charts showing gene expression changes in three groups of genes (909 genes losing 5mC, 753 genes gaining 5mC, and all genes).

In mammals, approximately 70% of genes are linked to promoter CpG islands that are constitutively hypomethylated, while the rest of promoters are depleted in CpGs and subject to context-dependent DNA methylation (Weber et al., 2007). Accordingly, we found that differentially methylated TSSs at the KIT^+^ to PS transition tend to overlap with low CpG promoters (Fig. 3B, right panel). In this context, TSSs tend to lose rather than gain 5mC levels. GO term analysis revealed that genes that lose 5mC at their TSS are enriched in genes involved in cell adhesion (presumably due to the changes in the *Pcdh* gene cluster seen earlier), sexual reproduction, spermatogenesis, and meiosis (Fig. 3C, Table S2). This is consistent with the demethylation and activation of late spermatogenesis genes described above. In contrast, genes that gain 5mC are enriched in immune-related functions and generally remain expressed at low levels during spermatogenesis (Fig. 3C and Fig. S2C). The low expression of these immune-related genes after germ cell differentiation may be related to the immune-privileged environment of the adluminal compartment established by the blood-testis barrier (Cheng & Mruk, 2012), although the underlying relationship remains to be determined.

We next examined the expression profiles of genes that lose 5mC. This group includes representative late spermatogenesis and meiotic genes, including *Spata16, Tdrd6, Slc22a16, Piwil1,* and *Spo11*, which are highly upregulated at the KIT^+^ to PS transition (Fig. 3D, Table S2). In contrast, several of the *Pcdh*γ genes show similar expression levels during spermatogenesis (Fig. 3D). Further, genes that lose 5mC levels tend to be upregulated at the KIT^+^ to PS transition compared to genes showing a gain of 5mC levels or all genes (Fig. 3E).

One special feature of spermatogenic gene expression is a specific expression of single exon genes. Indeed, certain CpG promoters of single exon genes are hypermethylated in soma but hypomethylated in late spermatogenesis (Kato & Nozaki, 2012). We found that, among 3,124 single-exon genes, 208 lose 5mC levels at the KIT^+^ to PS transition (Fig. S3A, Table S3). 62 of these genes are activated at the KIT^+^ to PS transition (Fig. S3B). Together, these data suggest that DNA demethylation at TSSs correlates with the activation of late spermatogenesis genes at the KIT^+^ to PS transition.

### DNA demethylation during spermatogenesis precedes the sites of nucleosome retention in spermatozoa

During spermatogenesis, histones are largely replaced with protamines. In spermatozoa, nucleosomes are retained mainly at hypomethylated TSSs, and these retained nucleosomes are implicated in paternal epigenetic inheritance (Erkek et al., 2013). Since we observed DNA demethylation at a number of TSSs at the KIT^+^ to PS transition, we hypothesized that such *de novo* hypomethylated TSSs might serve as the sites of nucleosome retention in spermatozoa. To test this hypothesis, we reanalyzed published MNase-seq data of sperm chromatin (Erkek et al., 2013). Specifically, we examined the degree of nucleosome retention (i.e., the enrichment of mononucleosomes) at various genomic regions detected in our MethylCap-seq data. Interestingly, we found a significant enrichment of mononucleosomes at promoters and exons that lost 5mC at the KIT^+^ to PS transition (termed KIT-unique methylated regions) (Fig. 4A). Consistent with these results, nucleosome retention was observed predominantly at promoters (49%) and exons (18%) in spermatozoa (Fig. 4B).

**Figure 4:**
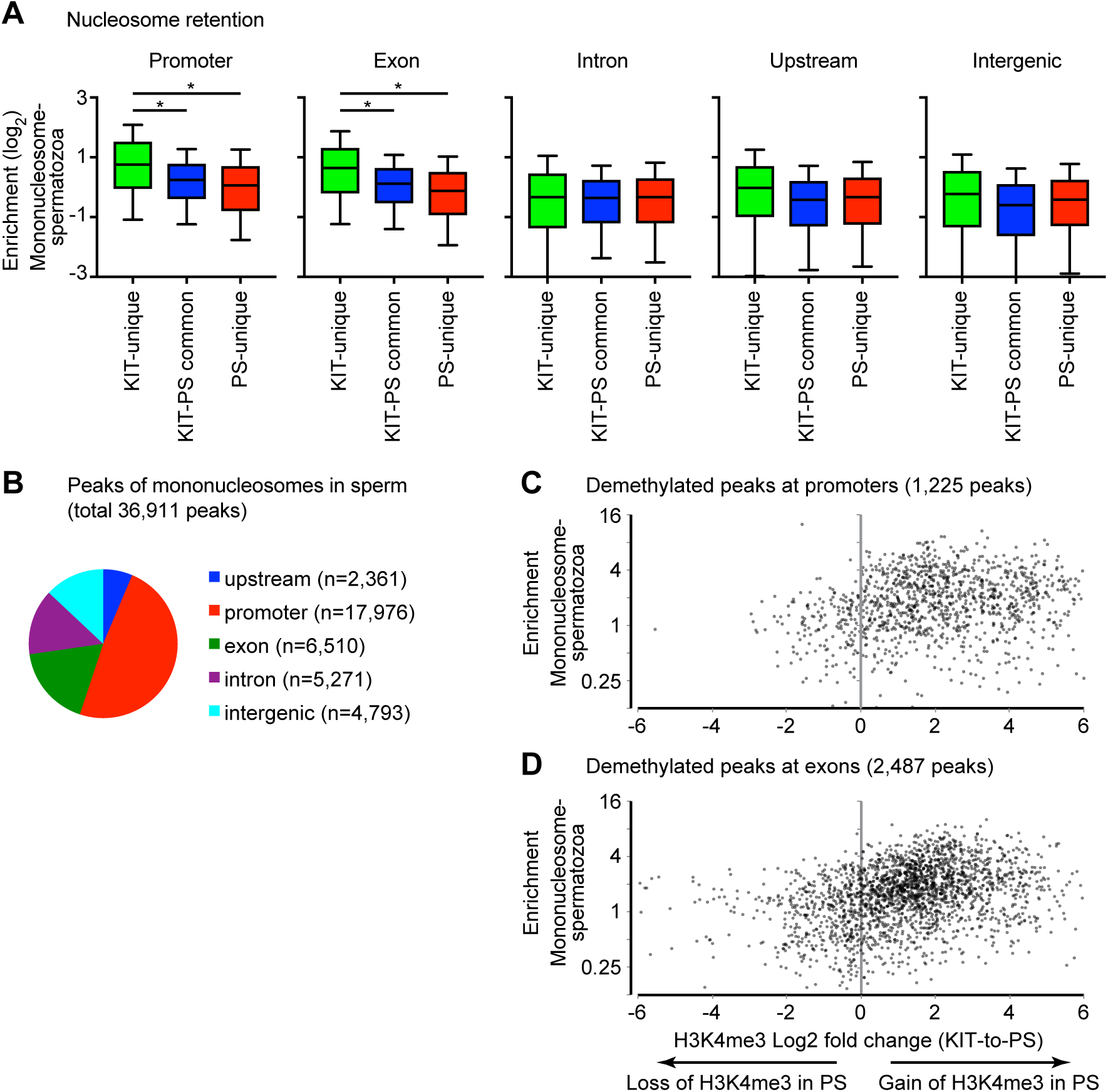
Sites of DNA demethylation precede sites of nucleosome retention during spermatogenesis. (A) Box-and-whisker plots showing mononucleosome enrichment at each genomic locus for each class of genomic regions. Central bars represent medians, the boxes encompass 50% of the data points, and the whiskers indicate 90% of the data points. * P-value < 0.0001, Mann–Whitney U test. (B) Genomic distribution of mononucleosome peaks in sperm (total 36,911 peaks): MNase-seq data (Erkek et al., 2013). Peaks are detected by MACS2 (Padj-value<0.01, Fold enrichment>5). (C, D) Changes in the enrichment of H3K4me3 (X-axis) and mononucleosomes in spermatozoa (Y-axes) at demethylated promoters (C) and exons (D) at the mitosis-to-meiosis transition.

To quantify the relationship between promoter DNA demethylation and nucleosome retention, we divided 1,225 promoter-associated demethylated peaks into four groups according to the magnitude of demethylation (Fig. S4). Sperm mononucleosome enrichment increased across groups with greater demethylation, with a significant ordered trend (Jonckheere–Terpstra test, *P* = 1.1 × 10⁻¹⁰). The magnitude of demethylation also showed a modest positive correlation with mononucleosome enrichment (Spearman’s ρ = 0.192, *P* = 1.31 × 10⁻¹¹). However, substantial variability remained among individual regions, indicating that promoter demethylation is associated with, but does not independently determine, nucleosome retention.

In spermatozoa, hypomethylated promoters are enriched with histones marked by H3K4me3, which counteracts DNA methylation (Erkek et al., 2013). Thus, we next examined whether promoters and exons that lose 5mC at the KIT^+^ to PS transition acquire H3K4me3. We reanalyzed our previous H3K4me3 ChIP-seq data of KIT^+^ spermatogonia and PS (Maezawa, Hasegawa, Alavattam, et al., 2018). Our scatter plot analysis shows that at demethylated promoters and exons, there is a gain of H3K4me3 at the KIT^+^ to PS transition, and these regions tend to show mononucleosome retention in spermatozoa (Fig. 4C, 4D). These results indicate that during spermatogenesis, DNA demethylation co-occurs with a gain of H3K4-methylation and precedes sites of nucleosome retention in sperm.

### DNA demethylation precedes nucleosome retention and bivalent chromatin in spermatozoa

Nucleosome retention sites in sperm are implicated in paternal epigenetic inheritance (Erkek et al., 2013). Accumulating evidence suggests incorporation of the histone variant H3.3 and enrichment of bivalent marks, i.e., active H3K4me3 and silent H3K27me3, at these sites (Erkek et al., 2013; Lismer & Kimmins, 2023; Sakashita et al., 2023). Thus, we sought to characterize H3.3 incorporation and bivalent marks of the sites that lose 5mC at the KIT^+^ to PS transition in spermatozoa. To this end, we classified the MethylCap-seq peaks of KIT^+^ spermatogonia into two classes: peaks which undergo demethylation from KIT^+^ spermatogonia to PS (Class I) and all other peaks (Class II) (Fig. 5A; Table S4). To determine whether the Class I regions overlap with previously identified TET1-dependent hypomethylated regions in sperm, we compared Class I regions with TET1-dependent sperm hypomethylated regions defined using whole-genome bisulfite sequencing data from *Tet1*-knockout sperm (Prasasya et al., 2024). Only limited overlap was observed between the two sets of regions (Fig. S5), indicating that most Class I regions do not correspond to the previously defined TET1-dependent sperm hypomethylated regions. In spermatozoa, Class I peaks showed greater aggregate mononucleosome and H3.3 enrichment than Class II peaks (Fig. 5A, B). Moreover, mononucleosome and H3.3 enrichment were strongly positively correlated across Class I peaks (Fig. 5C). These results suggest that spermatogenic demethylation precedes the incorporation of histone variant H3.3 and nucleosome retention in spermatozoa.

**Figure 5:**
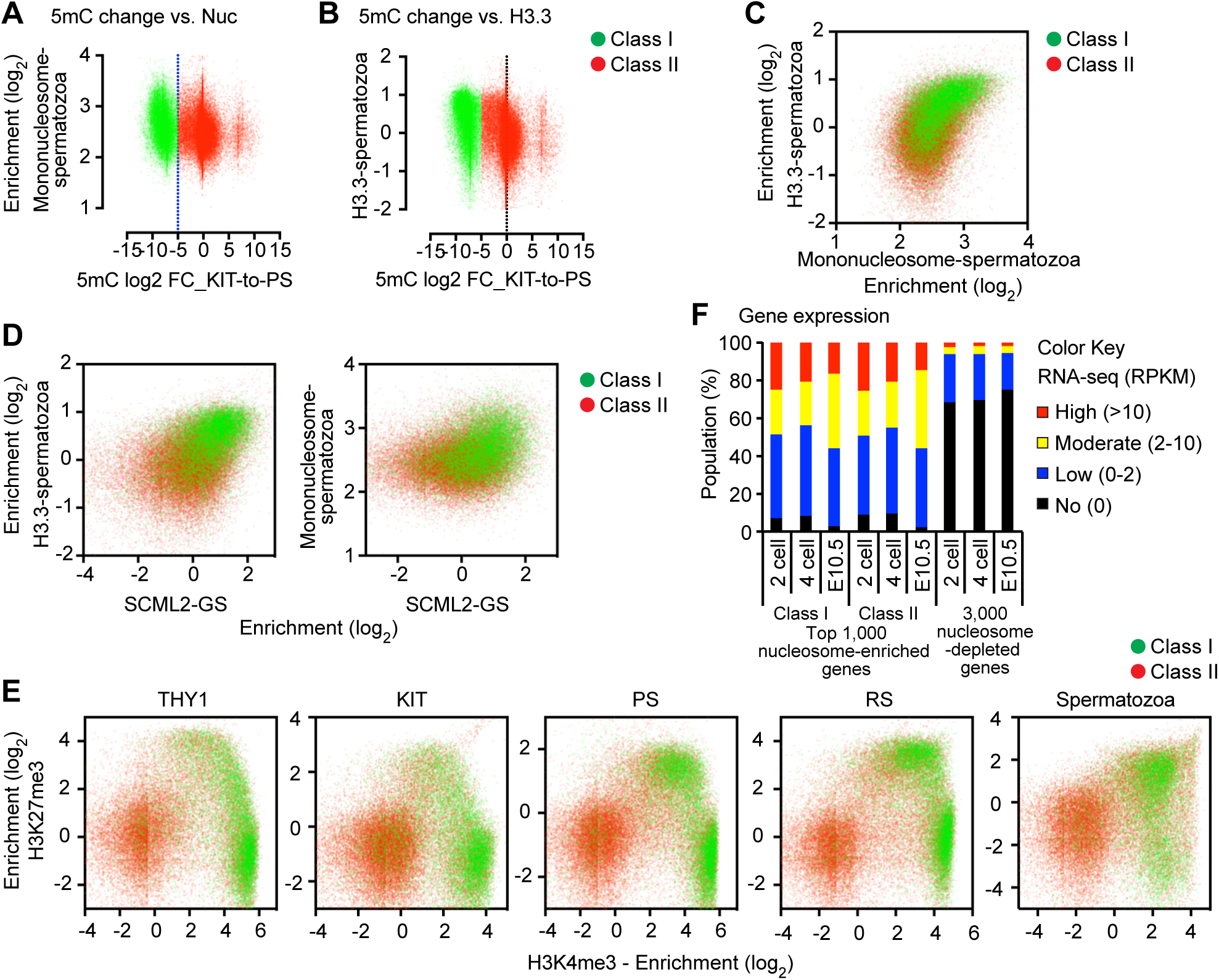
Sites of DNA demethylation precede sites of bivalent genomic domains in spermatozoa. (A) Mononucleosome enrichment in spermatozoa at genomic sites classified according to the magnitude of DNA demethylation from KIT+ spermatogonia to PS: Class I (green), log₂ fold change < −5; Class II (red), log₂ fold change ≥ −5. (B) H3.3 enrichment in the two classes of genomic sites in sperm. (C) Enrichment of H3.3 and nucleosomes (D) Enrichment of SCML2 in cultured germline stem (GS) cells, and H3.3 enrichment in sperm (left) and mononucleosome enrichment in sperm (right) (E) Enrichment of H3K27me3 and H3K4me3 during spermatogenesis and in spermatozoa. (F) RNA-seq analysis in embryos for three groups of genes. Top 1,000 nucleosome-enriched genes among class I and II peak-containing genes at promoters (7,577 and 7,261 genes) and bottom 3,000 nucleosome-enriched genes (Nucleosome-depleted genes).

We next wanted to examine whether bivalent marks are regulated at Class I regions during spermatogenesis. To this end, we focused on germline-specific Polycomb protein SCML2, which has a high affinity to hypomethylated DNA and is a critical regulator of germline transcriptomes and bivalent chromatin (Hasegawa et al., 2015; Maezawa, Hasegawa, Alavattam, et al., 2018; Maezawa et al., 2020; Maezawa, Yukawa, et al., 2018). SCML2 binding sites in spermatogonia predict the sites of H3K27me3 deposition in late spermatogenesis, and loss of SCML2 results in H3K27me3 depletion and defective bivalent domains (Maezawa, Hasegawa, Yukawa, et al., 2018), leading to abnormal paternal epigenetic inheritance (Sakashita et al., 2023). We, therefore, examined the enrichment of SCML2 by reanalyzing our previous SCML2 ChIP-seq data (Hasegawa et al., 2015). The Class I peaks tend to show a higher degree of SCML2 and H3.3 enrichment as well as a higher degree of nucleosome retention than Class II peaks (Fig. 5D). Because SCML2 binds to hypomethylated promoters enriched with H3K4me3 and establishes H3K27me3 in PS (Maezawa, Hasegawa, Alavattam, et al., 2018), we next examined enrichment of H3K4me3 and H3K27me3 at Class I and II peaks during spermatogenesis. In THY1^+^ and KIT^+^ spermatogonia, Class I peaks are enriched with H3K4me3. In contrast, Class II peaks are not enriched with H3K4me3 (Fig. 5E). The Class I peaks gradually gain H3K27me3 during differentiation from PS to spermatozoa, and the majority are enriched for both H3K4me3 and H3K27me3 in spermatozoa (Fig. 5E). Taken together, we conclude that spermatogenic demethylation precedes nucleosome retention enriched with bivalent marks H3K4me3 and H3K27me3 in spermatozoa.

Finally, we examined the relationship between spermatogenic demethylation-associated regions, sperm nucleosome enrichment, and embryonic gene expression. Genes whose promoters are enriched with nucleosomes in spermatozoa (top 1,000 nucleosome-enriched genes among class I and II peak-containing genes at promoters) are highly expressed in embryogenesis at 2-cell and 4-cell stages and at embryonic day 10.5 (E10.5) (Fig. 5F), although we did not observe an apparent difference between these two groups. This enrichment was greater than that observed for the bottom 3,000 genes ranked by promoter nucleosome enrichment in spermatozoa. Thus, nucleosome enrichment in spermatozoa is associated with gene expression in early embryos.

These results suggest that the genes that undergo spermatogenic demethylation are enriched with nucleosomes in spermatozoa and are expressed in embryos of the next generation.

## Discussion

In this study, we have mapped site-specific DNA methylation during the transition from mitotic spermatogonia to meiotic spermatocytes. We observed that promoters of a large number of late spermatogenesis genes undergo DNA demethylation at the KIT^+^ spermatogonia to PS transition, indicating that DNA demethylation might mark late spermatogenesis genes for induction. Our data also suggest that these demethylated regions precede sites of nucleosome retention in spermatozoa. Specifically, we propose that demethylated regions in meiotic spermatocytes acquire H3K4me3 and subsequently acquire SCML2-dependent H3K27me3, thereby establishing bivalent marks at hypomethylated nucleosome retention sites in spermatozoa (Fig. 6). Previous studies showed that SCML2 is highly expressed in undifferentiated spermatogonia and is associated with hypomethylated promoters at this stage, whereas PRC2-mediated H3K27me3 accumulates predominantly after meiotic entry (Hasegawa et al., 2015; Maezawa, Hasegawa, Yukawa, et al., 2018). Thus, SCML2 occupancy precedes H3K27me3 deposition and may mark regions that subsequently acquire Polycomb-mediated chromatin states during spermatogenesis. In support of this model, we recently demonstrated that SCML2 mediates paternal epigenetic inheritance through sperm chromatin (Sakashita et al., 2023).

**Figure 6:**
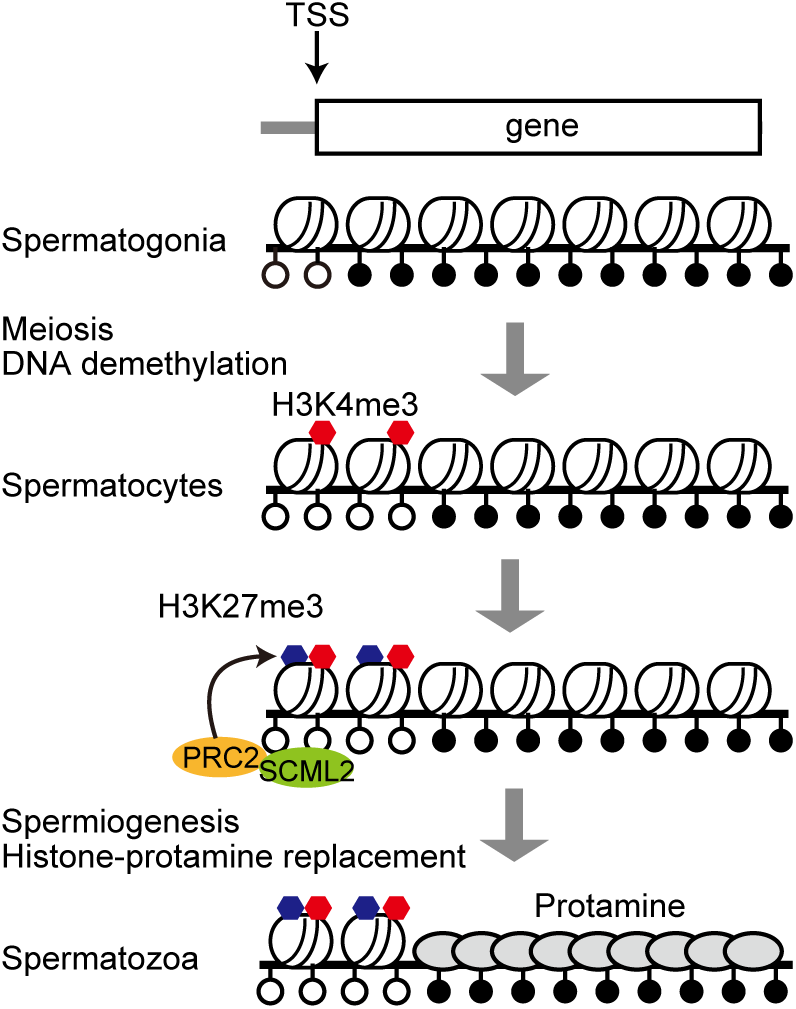
Model: Sites of DNA demethylation precede sites of nucleosome retention during spermatogenesis. Genomic regions that are demethylated during the transition from mitotic spermatogonia to meiotic spermatocytes acquire H3K4me3 in meiotic spermatocytes, leading to SCML2-mediated deposition of H3K27me3, thereby establishing persisting bivalent marks at these hypomethylated nucleosome retention sites in spermatozoa.

A recent study demonstrated that experimentally induced DNA hypomethylation in the male germline increases nucleosome retention in mature sperm, particularly at CpG-rich regions (Fanourgakis et al., 2025). The same study further showed that reduced sperm DNA methylation renders paternal alleles permissive for H3K4me3 establishment in early embryos, independently of the direct inheritance of sperm-borne H3K4me3. Consistent with the findings of Fanourgakis et al., our data show that genomic regions undergoing demethylation during spermatogenesis are preferentially associated with nucleosome retention in mature sperm. Our study further identifies when these hypomethylated regions emerge during normal spermatogenesis, suggesting that site-specific demethylation during the mitosis-to-meiosis transition contributes to the establishment of sperm epigenetic states associated with nucleosome retention and subsequent embryonic chromatin regulation. The extent of promoter demethylation showed a modest positive association with sperm nucleosome enrichment; however, substantial variability among individual regions indicates that DNA demethylation alone is insufficient to determine nucleosome retention. Additional features, including histone variants, histone modifications, CpG density, and the local chromatin environment, are therefore likely to contribute to the establishment of retained nucleosomes during spermiogenesis.

Our data reveal a previously unrecognized feature of the male germline: site-specific DNA demethylation during meiotic progression is linked to sperm chromatin features implicated in paternal epigenetic inheritance. We propose that this meiotic DNA demethylation contributes to the establishment of sperm epigenetic states associated with gene regulation in the embryo. In 1984, Robin Holliday proposed that an aspect of the biological significance of meiosis is the reprogramming of gametes for fertilization (Holliday, 1984), and our data provide a foundation for this concept.

The localized gains of DNA methylation detected by MethylCap-seq are not necessarily inconsistent with the global reduction in 5mC observed by ELISA and immunostaining, because these approaches measure distinct aspects of DNA methylation. ELISA and immunostaining reflect overall genomic 5mC abundance, whereas MethylCap-seq detects regional enrichment of methylated DNA primarily within uniquely mappable genomic regions. Thus, localized gains and losses of DNA methylation can occur despite an overall reduction in genomic 5mC. In addition, methylation changes within highly repetitive regions are not fully captured by our MethylCap-seq analysis. Therefore, the global and locus-specific changes observed here likely represent complementary aspects of DNA methylation remodeling during spermatogenesis.

The site-specific DNA methylation changes described in this study may arise through mechanisms distinct from the transient hemimethylation that occurs during premeiotic S phase (Gaysinskaya et al., 2018; Huang et al., 2023). This transient state results from delayed maintenance methylation following premeiotic DNA replication and is therefore global rather than site-specific.

Recent work demonstrated that TET1 is required for the establishment of a subset of hypomethylated regions in mature sperm (Prasasya et al., 2024). However, the regions demethylated during the KIT-positive spermatogonia-to-pachytene spermatocyte transition showed only limited overlap with these TET1-dependent sperm hypomethylated regions.

Although differences in developmental stage, assay, and region definition should be considered, this finding suggests that much of the site-specific demethylation identified here may occur independently of TET1. Other TET family members may contribute to this process. Alternatively, some of these changes may be related to incomplete maintenance methylation during premeiotic DNA replication. Further studies will be required to define the underlying mechanisms.

The paternal genome undergoes extensive reprogramming after fertilization. For example, paternal H3.3 on promoters disappears after fertilization (Ishiuchi et al., 2021), H3K27me3 is reprogrammed during preimplantation development (Zheng et al., 2016) and the paternal genome undergoes passive and active DNA demethylation (Guo et al., 2014; Shen et al., 2014; Wang et al., 2014). Thus, a major unsolved mystery is how paternal epigenetic states are inherited and maintained after fertilization. One possibility is that the paternal epigenetic states, reflecting the memory of paternal hypomethylated DNA, escape epigenetic reprogramming and persist at the promoter regions of the target genes throughout germ cell development.

Nevertheless, this possibility has been confounded by the debates about the sites of nucleosome retention in spermatozoa (Yin et al., 2023). Therefore, an important next step is to elucidate how the paternal hypomethylated DNA regions are regulated in the sperm-to-zygote transition.

## Materials and Methods

### Animals

Mice on the C57BL/6 background were maintained and used according to the guidelines of the Institutional Animal Care and Use Committee (protocol no. IACUC2018–0040) at Cincinnati Children’s Hospital Medical Center.

### Germ cell fractionation

Wild-type mice on the C57BL/6 background (at least 12 independent mice at 8-12 weeks of age or at least 20 independent mice at 6-8 days of age) were used for isolation of germ cells.

Pachytene spermatocytes and round spermatids were isolated via BSA gravity sedimentation as previously described (Bellve, 1993). Purity was confirmed by nuclear staining with Hoechst 33342 using fluorescence microscopy. In keeping with previous studies from the Namekawa lab (Alavattam et al., 2019; Hasegawa et al., 2015; Maezawa, Hasegawa, Yukawa, et al., 2018; Maezawa, Yukawa, et al., 2018), only fractions with a mean purity of ≥90% were used to extract genomic DNA.

Spermatogonia were isolated as described previously (Hammoud et al., 2014) and collected from C57BL/6 wild-type mice aged 6–8 days. Testes were collected in a 24-well plate in Dulbecco’s Modified Eagle Medium (DMEM) supplemented with GlutaMax (Thermo Fisher Scientific), non-essential amino acids (NEAA) (Thermo Fisher Scientific), and penicillin and streptomycin (Thermo Fisher Scientific). After removing the *tunica albuginea* membrane, testes were digested with collagenase (1 mg/ml) at 34°C for 20 min to remove interstitial cells, then centrifuged at 188×*g* for 5 min. Tubules were washed with the medium and then digested with trypsin (2.5 mg/ml) at 34°C for 20 min to obtain a single-cell suspension. Cells were filtered with a 40-μm strainer to remove Sertoli cells, and the cell suspension was plated in a 24-well plate for 1 h in the medium supplemented with 10% fetal bovine serum, which promotes adhesion of remaining somatic cells. Cells were washed with magnetic cell-sorting (MACS) buffer (PBS supplemented with 0.5% BSA and 5 mM EDTA) and incubated with CD117 (KIT) MicroBeads (Miltenyi Biotec) on ice for 20 min. Cells were washed and resuspended with MACS buffer and filtered with a 40-μm strainer. Cells were separated by autoMACS Pro Separator (Miltenyi Biotec) with the program “possel.” Cells in the flow-through fraction were washed with MACS buffer and incubated with CD90.2 (THY1) MicroBeads (Miltenyi Biotec) on ice for 20 min. Cells were washed and resuspended with MACS buffer and filtered with a 40-μm strainer. Cells were separated by autoMACS Pro Separator (Miltenyi Biotec) with the program “posseld.” Purity was confirmed by immunostaining.

### Quantification of 5mC

Genomic DNA was purified from 1–10 × 10⁵ isolated spermatogonia, spermatocytes, or spermatids using the Genomic DNA Clean & Concentrator Kit (Zymo Research) according to the manufacturer’s instructions. Global 5mC levels were quantified using 100–130 ng of genomic DNA with the MethylFlash™ Methylated 5mC DNA Quantification Kit (Colorimetric) (Epigentek) according to the manufacturer’s instructions.

Biological replicates consisted of six independent samples for THY1-positive and KIT-positive spermatogonia and five independent samples for pachytene spermatocytes and round spermatids. Data are presented as mean ± SEM. Statistical significance was assessed by one-way ANOVA followed by Tukey’s multiple-comparison test. Differences were considered significant at P < 0.05.

### Histological analysis and immunofluorescence

For the preparation of testicular paraffin blocks, testes were fixed with 4% paraformaldehyde (PFA) overnight at 4°C with gentle inverting. Testes were dehydrated and embedded in paraffin. For immunofluorescence analysis of testicular sections, antigen retrieval was performed by boiling the slides in Target Retrieval Solution (DAKO) for 10 min and letting the solution cool for 30 min. For the detection of 5mC, sections were incubated in 4 N HCl for 1 h at room temperature and then neutralized in 100 mM Tris-HCl (pH 8.5) for 10 min, followed by a standard immunostaining protocol. Sections were blocked with Blocking One Histo (Nacalai) for 1 h at room temperature and then incubated with anti-5mC (1/200 dilution, Eurogentec, clone 33D3), anti-ZBTB16 (1/200 dilution, Santa Cruz, sc22839), anti-H1T (1/1000 dilution, gift from Dr. Mary Ann Handel), goat anti-KIT (1/200 dilution, R&D Systems, AF1356), rabbit anti- STRA8 (1/500 dilution, Abcam, ab49602), goat anti-DMC1 (1/200 dilution, Santa Cruz, sc8973), rabbit anti-γH2AX (1/500 dilution, Abcam, ab11174), or mouse anti-γH2AX (1/500 dilution, Merck Millipore, 05-636) antibodies overnight at 4°C. The resulting signals were detected by incubation with secondary antibodies conjugated to Alexa Fluor dyes (Thermo Fisher Scientific or Jackson ImmunoResearch). Sections were counterstained with DAPI. Images were obtained via a laser scanning confocal microscope A1R (Nikon) and processed with NIS-Elements (Nikon) and ImageJ (National Institutes of Health) (Schneider et al., 2012).

### MethylCap-seq

Genomic DNA was fragmented by sonication and methylated DNA fragments were captured by MethylAffinity Methylated DNA Enrichment Kit (GeneCopoeia). DNA libraries were created by ChIPmentation method (Schmidl et al., 2015). Briefly, methylated DNA fragment-bound beads were resuspended in 30 μl of the tagmentation reaction buffer (10 mM Tris-HCl pH 8.0 and 5 mM MgCl_2_) containing 1 μl Tagment DNA Enzyme from the Nextera DNA Sample Prep Kit (Illumina) and incubated at 37°C for 10 min in a thermal cycler. The beads were washed twice with 150 μl cold wash buffer (50 mM Tris-HCl pH 8.0, 150 mM NaCl, 1 mM EDTA, 0.1% SDS, 0.1% NaDOC, and 1% Triton X-100), incubated with elution buffer (10 mM Tris-HCl pH 8.0, 1 mM EDTA, 250 mM NaCl, 0.3% SDS, 0.1 μg/μl Proteinase K) at 42°C for 30 min. DNA was purified with the MinElute Reaction Cleanup Kit (Qiagen) and amplified with NEBNext High-Fidelity 2× PCR Master Mix (NEB). Amplified DNA was purified by Agencourt AMPure XP (Beckman Coulter). Afterward, DNA fragments in the 250- to 500-bp size range were prepared by agarose gel size selection. DNA libraries were adjusted to 5 nM in 10 mM Tris-HCl pH 8.0 and sequenced with an Illumina HiSeq 2500.

### Data analysis

Data analysis for MethylCap-seq, ChIP-seq, and RNA-seq was performed in the BioWardrobe Experiment Management System (Kartashov & Barski, 2015). For MethylCap-seq and ChIP-seq analysis, reads were aligned to mouse genome mm10 with Bowtie (version 1.2.0, (Langmead et al., 2009)), assigned to NCBI RefSeq genes or isoforms, and coverage was displayed on a local mirror of the UCSC genome browser (Meyer et al., 2013). MethylCap-seq peaks were identified using MACS2 (version 2.1.1.20160309, (Zhang et al., 2008)) with a default parameter setting for narrow peak detection in BioWardrobe. Pearson correlations for the genome-wide enrichment of the peaks among MethylCap-seq library replicates were analyzed using SeqMonk (Babraham Institute).

MAnorm, software designed for quantitative comparisons of ChIP-seq data sets (Shao et al., 2012), was used to compare the genome-wide MethylCap-seq peaks among stages in spermatogenesis. Unique peaks were defined using the following criteria: (1) defined as “unique” by the MAnorm algorithm; (2) P-value <0.01; (3) raw counts of unique reads >10. Peaks common to two stages were defined using the following criteria: (1) defined as “common” by MAnorm algorithm; (2) raw read counts of both stages >10.

Average tag density profiles were calculated around transcription start sites for gene sets of somatic/progenitor genes, late spermatogenesis genes, constitutive active genes, and constitutive inactive genes, as described previously (Sin et al., 2015). The resulting graphs were smoothed in 200-bp windows. Enrichment levels for MethylCap-seq experiments were calculated for 10-kb windows, promoter regions of genes (±5 kb surrounding TSSs).

Enrichment levels for MethylCap-seq and ChIP-seq experiments were calculated for 4-kb windows, promoter regions of genes (±2 kb surrounding TSSs). To normalize tag values, read counts were multiplied by 1,000,000 and then divided by the total number of reads in each nucleotide position. The total amount of tag values in promoter or enhancer regions was calculated as enrichment.

To detect differentially methylated regions around TSSs among stages in spermatogenesis, a read count output file was input to the DESeq2 package (version 1.16.1); then, the program functions DESeqDataSetFromMatrix and DESeq were used to compare each MethylCap-seq enrichment between two biological samples. Differentially methylated regions were identified through binominal tests, thresholding Benjamini-Hochberg- adjusted *P* values to <0.05. To perform gene ontology analyses, the functional annotation clustering tool in DAVID (version 6.8) was used, and a background of all mouse genes was applied.

For RNA-seq analysis, reads were aligned by STAR (version STAR_2.5.3a) (Dobin et al., 2013) with default arguments except --outFilterMultimapNmax 1 and --outFilterMismatchNmax 2. The --outFilterMultimapNmax parameter was used to allow unique alignments only, and the -- outFilterMismatchNmax parameter was used to allow a maximum of 2 errors. NCBI RefSeq annotation from the mm10 UCSC genome browser was used, and canonical TSSs (1 TSS per gene) were analyzed. All reads from the resulting .bam files were split for related isoforms with respect to RefSeq annotation.

For the analysis shown in Figure S4, promoter-associated demethylated peaks were ranked according to the magnitude of DNA demethylation and divided into four equal-sized groups. Ordered differences in sperm mononucleosome enrichment across the four groups were assessed using the Jonckheere–Terpstra test. The association between the magnitude of DNA demethylation and sperm mononucleosome enrichment values was evaluated using Spearman’s rank correlation.

### Comparison with *Tet1*-deficient sperm methylation datasets

Genomic regions undergoing DNA methylation changes in Tet1-deficient sperm were obtained from the processed data accompanying Prasasya et al. (2024). Hyper- and hypomethylated regions identified in *Tet1*-knockout sperm were compared with the Class I regions identified in this study using BEDTools intersect (version 2.31.0) with default parameters. Genomic overlaps were evaluated based on the mm10 (GRCm38) mouse genome assembly.

## Data availability

NGS datasets used in this study are publicly available and referenced within the article. All the MethylCap-seq data generated in this study are deposited to the Gene Expression Omnibus (GEO) under accession code GSE262669.

## Author Contributions

S.M. and S.H.N. designed research; S.M. performed research; S.M., M.Y., A.S., Y.Y., T.S., A.B., and S.H.N. analyzed data; and S.M. and S.H.N. wrote the paper. S.H.N. supervised the project.

## Acknowledgments

We thank members of the Namekawa and Maezawa labs for discussion and helpful comments regarding the manuscript.

---

**Figure S1. Changes in 5mC levels during spermatogenesis.**

(A, B) Immunostaining of adult testis sections for 5mC and the indicated stage-specific markers. Differentiating spermatogonia and preleptotene spermatocytes are identified by KIT and STRA8, respectively (A). Leptotene and pachytene spermatocytes are identified by DMC1 and γH2AX, respectively (B). Boxed regions are magnified in the right panels. Scale bars, 50 μm in the overview images and 5 μm in the magnified images.

**Figure S2. Locus-specific DNA methylation changes during spermatogenesis.**

(A) Genome-browser tracks showing MethylCap-seq and whole-genome bisulfite sequencing (WGBS) data at the *Spata16* and *Slc22a16* loci during spermatogenesis.

(B) Genome-browser tracks showing MethylCap-seq, WGBS, and H3K4me3 ChIP-seq data at the *Pcdhγ* locus during spermatogenesis.

(C) RNA-seq heatmap of 753 genes gaining 5mC around their TSSs from KIT+ spermatogonia to pachytene spermatocytes.

**Figure S3. DNA methylation and expression of single-exon genes during spermatogenesis.**

(A) Overlap of single-exon genes with genes losing or gaining 5mC around their TSSs from KIT^+^ spermatogonia to pachytene spermatocytes. The 208 single-exon genes losing 5mC include 33 olfactory receptor genes, 14 *Pcdhb* genes, and 16 miRNA genes.

(B) Numbers of genes activated or repressed by at least fourfold from KIT⁺ spermatogonia to pachytene spermatocytes among single-exon genes and all genes with differentially methylated TSSs. RPKM > 5 was required in pachytene spermatocytes for activated genes and in KIT⁺ spermatogonia for repressed genes. SEG, single-exon genes.

**Figure S4: Promoter DNA demethylation is associated with increased nucleosome enrichment.**

Promoter-associated demethylated peaks (n = 1,225) were ranked by the magnitude of DNA demethylation and divided into four equal-sized groups. Box plots show the median, interquartile range, and values within 1.5 times the interquartile range. The ordered trend across the four groups was evaluated using the Jonckheere–Terpstra test. The association between the magnitude of DNA demethylation and sperm mononucleosome enrichment was assessed using Spearman’s rank correlation.

**Figure S5. Limited overlap between Class I regions and DNA methylation changes in Tet1- knockout sperm.** Venn diagrams show the overlap of Class I regions identified in this study with regions reported to be hypermethylated or hypomethylated in *Tet1*-knockout sperm based on processed whole-genome bisulfite sequencing data from Prasasya et al. (2024). The numbers indicate the number of regions in each category and their intersections.

**Table S1. Common and transition-specific MethylCap-seq peaks identified across all spermatogenic transitions (Fig. 2A)**

**Table S2. Differentially methylated transcription start sites and associated gene expression data (Fig. 3B–D)**

**Table S3. Single-exon genes and associated gene expression data (Fig. S3A, B)**

**Table S4. Class I and Class II loci defined by the extent of DNA demethylation (Fig. 5)**

