## Supplemental Figure S1-5 for "Site-specific DNA demethylation during spermatogenesis precedes nucleosome retention in mouse sperm"

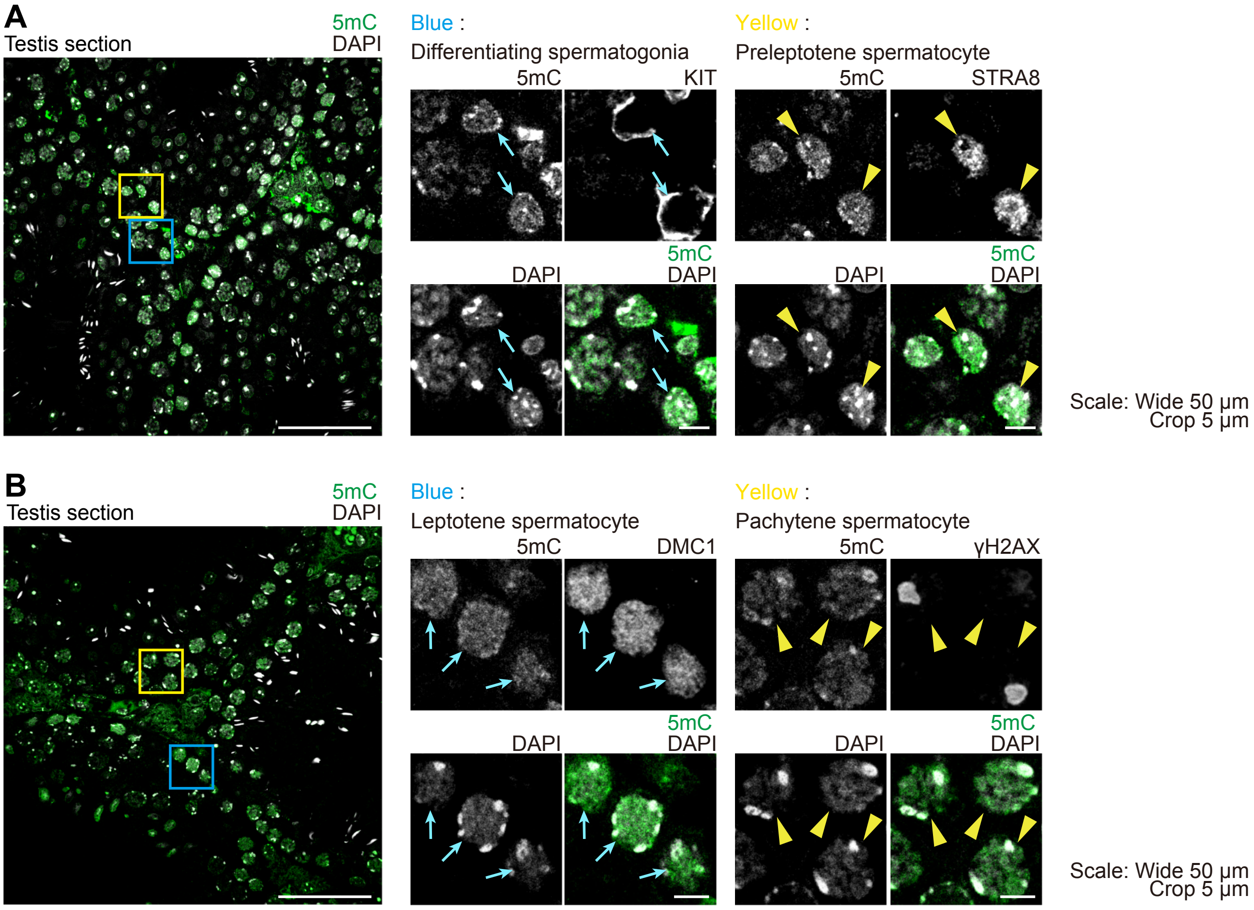
A
Testis section
5mC
DAPI
Blue :
Differentiating spermatogonia
5mC
KIT
DAPI
5mC
DAPI
Yellow :
Preleptotene spermatocyte
5mC
STRA8
DAPI
5mC
DAPI
Scale: Wide 50 µm
Crop 5 µm
B
Testis section
5mC
DAPI
Blue :
Leptotene spermatocyte
5mC
DMC1
DAPI
5mC
DAPI
Yellow :
Pachytene spermatocyte
5mC
γH2AX
DAPI
5mC
DAPI
Scale: Wide 50 µm
Crop 5 µm

Maezawa et al. Figure S2

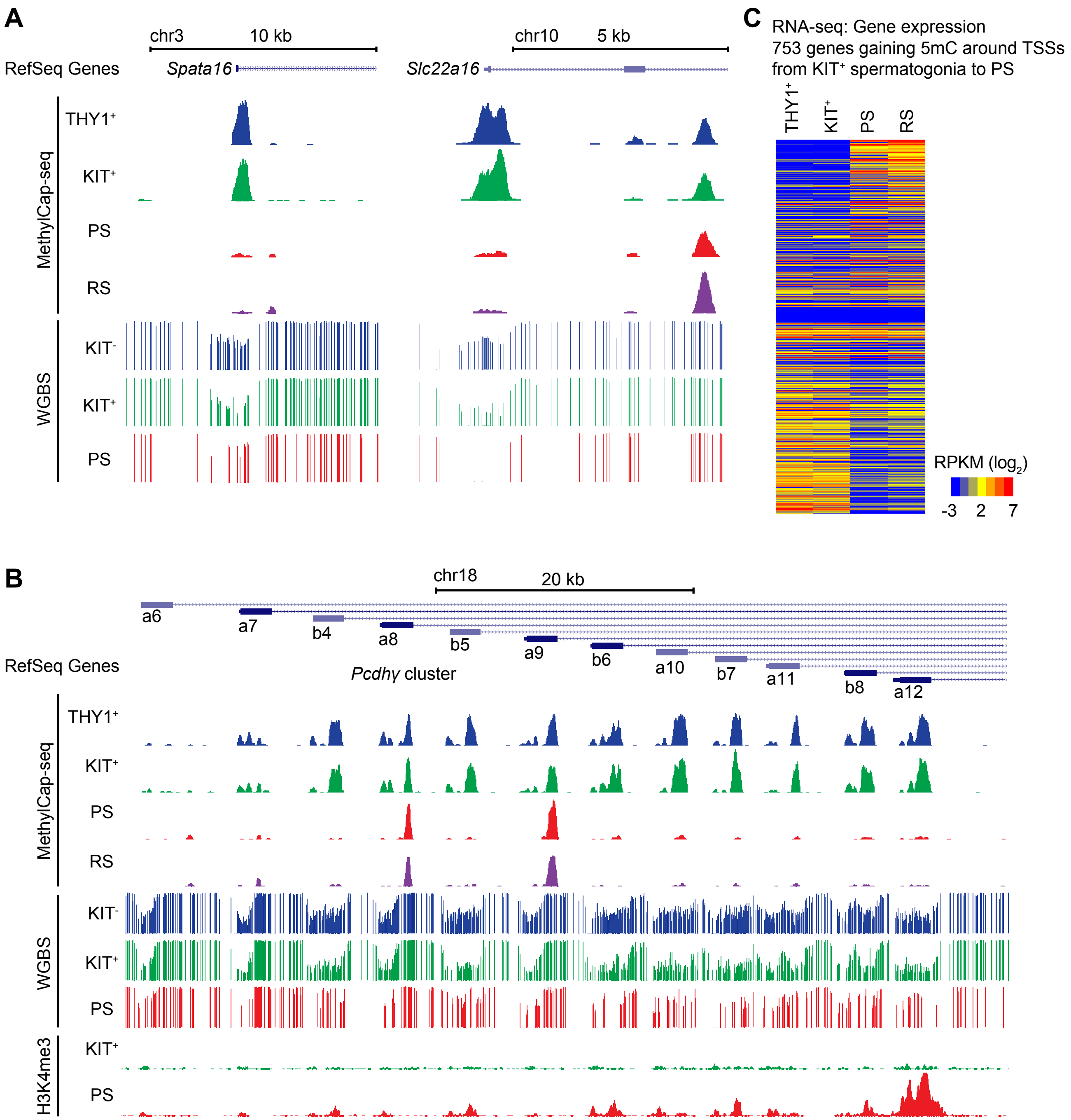

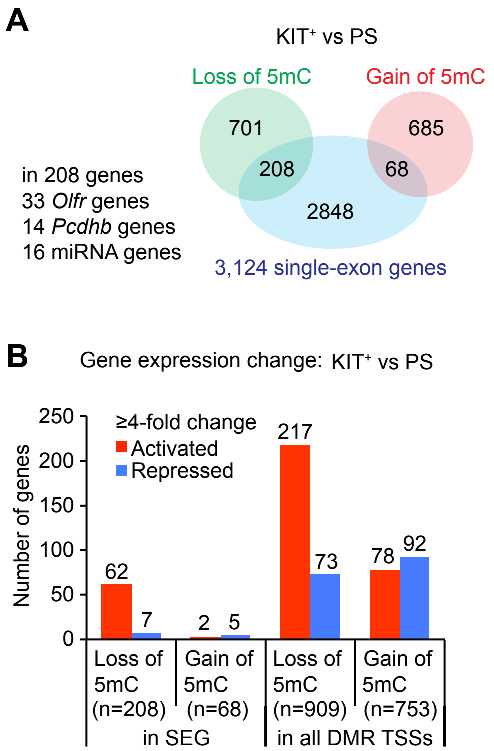
A
KIT+ vs PS
Loss of 5mC
Gain of 5mC
701
685
208
68
2848
in 208 genes
33 Olfr genes
14 Pcdhb genes
16 miRNA genes
3,124 single-exon genes
B
Gene expression change: KIT+ vs PS
≥4-fold change
Activated
Repressed
Number of genes
250
200
150
100
50
0
62
7
2
5
217
73
78
92
Loss of 5mC (n=208)
Gain of 5mC (n=68)
in SEG
Loss of 5mC (n=909)
Gain of 5mC (n=753)
in all DMR TSSs

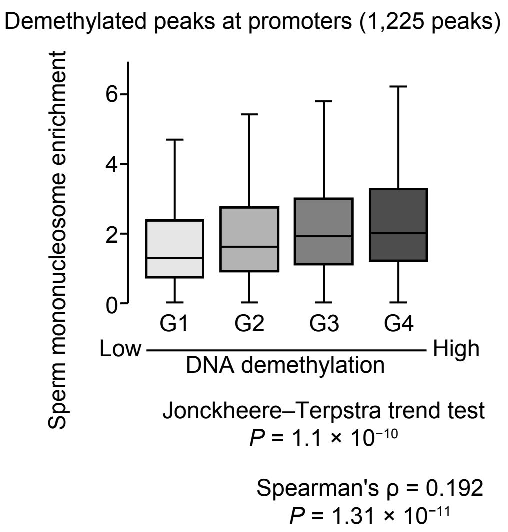
Demethylated peaks at promoters (1,225 peaks)
Sperm mononucleosome enrichment
6
4
2
0
G1
G2
G3
G4
Low
High
DNA demethylation
Jonckheere–Terpstra trend test
P = 1.1 × 10⁻¹⁰
Spearman's ρ = 0.192
P = 1.31 × 10⁻¹¹

Maezawa et al. Figure S5

Overlap of Class I regions with TET1-dependent sperm DMRs

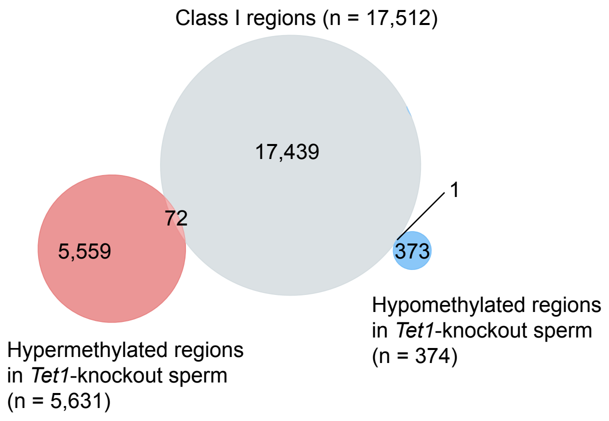
